# Two cases of type-a *Haemophilus influenzae* meningitis within the same week in the same hospital are phylogenetically unrelated but recently exchanged capsule genes

**DOI:** 10.1101/619502

**Authors:** Yves Terrat, Lauge Farnaes, John Bradley, Nicolas Tromas, B. Jesse Shapiro

## Abstract

*H. influenzae* causes common and sometimes severe pediatric disease including chronic obstructive respiratory disease, otitis media, and infections of the central nervous system. Serotype b strains, with a b-type capsule, have been the historical cause of invasive disease, and the introduction of a serotype b-specific vaccine has led to their decline. However, unencapsulated or non-b-type *H. influenzae* infections are not prevented by the vaccine and appear to be increasing in frequency. Here we report two pediatric cases of severe central nervous system *H. influenzae* infection presenting to the same hospital in San Diego, California during the same week in January 2016. Due to good vaccine coverage in this part of the world, *H. influenzae* cases are normally rare and seeing two cases in the same week was unexpected. We thus suspected a recent transmission chain, and possible local outbreak. To test this hypothesis, we isolated and sequenced whole genomes from each patient and placed them in a phylogenetic tree spanning the known diversity of *H. influenzae*. Surprisingly, we found that the two isolates (H1 and H2) belonged to distantly related lineages, suggesting two independent transmission events and ruling out a local outbreak. Despite being distantly related, H1 and H2 belong to two different lineages that appear to engage in frequent horizontal gene transfer (HGT), suggesting overlapping ecological niches. Together, our comparative genomic analysis supports a scenario in which an f-type ancestor of H2 arrived in North America around 2011 and acquired an a-type capsule by recombination (HGT) with a recent ancestor of H1. Therefore, as in other bacterial pathogens, capsule switching by HGT may be an important evolutionary mechanism of vaccine evasion in *H. influenzae*.

**OUTCOME:** Two cases of severe central nervous system *H. influenzae* infection occurred during the same week in the same hospital in San Diego, California – a region where such infections are usually very rare due to vaccine coverage. We thus suspected a local outbreak of an *H. influenzae* clone not covered by the vaccine. Using whole genome sequencing and phylogenetic analysis of two isolates (H1 and H2, one from each patient), we found that they were distantly related, rapidly ruling out a local outbreak and suggesting independent transmission events. This result highlights the potential for rapid global spread of non-vaccine *H. influenzae* strains. In this case, both H1 and H2 both encoded a-type capsules, whereas the vaccine targets b-type capsules. We also present comparative genomic evidence that a recent f-type ancestor of H2 acquired an a-type capsule locus from a recent ancestor of H1, and that this horizontal gene transfer (HGT) event likely happened in the past decade in North America, but probably not in the San Diego hospital. These results highlight the potential importance of HGT in the capsule locus in allowing *H. influenzae* to escape vaccine coverage.

**DATA SUMMARY:** *H. influenzae* H1 and H2 genome sequences have been deposited in NCBI under BioProject PRJNA534512.

**CONFLICT OF INTEREST STATEMENT:** The authors declare that they have no conflict of interest to report.

## INTRODUCTION

*Haemophilus influenzae* is traditionally classified into encapsulated or unencapsulated strains, with encapsulated strains being subdivided into serotypes (or types) a-f, each with a distinct type of polysaccharide capsule. Type-b has long been associated with invasive disease (Pittman, 1931) and has thus been a major vaccine target. Since the introduction of vaccine against type-b *H. influenzae*, a dramatic decrease of severe cases has been observed (Peltola, 2000). However, this drop in severe type-b infections was followed by an increase of acute infections caused by non-b-type (*i.e*. a, c, d, e, and f capsule types) and non-typeable (unencapsulated) strains (Ladhani et al., 2012; Headrick et al., 2018; Tsang et Ulanova, 2017; Giufrè et al 2017).

As a common surface antigen and vaccine target, the capsule is a target of diversifying selection and the capsule locus is a hotspot of recombination in bacterial pathogens including *Klebsiella pneumoniae* (Wyres et al., 2015), *Streptococcus pneumoniae* (Mostowy et al., 2017), and *Neisseria meningitidis* (Bartley et al., 2017) – but has been less thoroughly studied in *H. influenzae*. Many (but not all) natural *H. influenzae* isolates are competent for DNA uptake (Maughan et Redfield, 2009), and *H. influenzae* housekeeping and lipopolysaccharide genes are inferred to undergo relatively frequent recombination (Vos et Didelot, 2009). Capsule types tend to be associated with particular phylogenetic lineages of *H. influenzae*, leading to the assertion that capsule genes evolve clonally, with limited recombination (De Chiara et al., 2014). On the other hand, the capsule locus can be deleted naturally by recombination (Kroll et al., 1988, and the capsule locus is occassionally recombined among phylogenetically distant lineages (Musser et al., 1988). Thus, capsular recombination in *H. influenzae* appears to be relatively rare, but its impact on the evolution and epidemiology of *H. influenzae* infections could be substantial. For example, capsular switching could allow successful pathogenic lineages to evade vaccines and persist. Alternatively, if capsular recombination is limited, we would expect vaccine lineages to be replaced with other lineages, encoding different capsules.

Here we describe two *H. influenzae* genomes, each isolated from a meningitis patient at Rady Children’s Hospital (California, USA) within one week of each other in January of 2016. As such severe central nervous system infections are extremely rare in Western countries since the introduction of *H. influenzae* vaccines in the 1990s, the appearance of two cases in a such narrow geographic and time window lead us to address the following questions using a comparative genomic approach:

1. Are the two strains closely related, suggesting an outbreak of a particular *H. influenzae* clone – possibly a vaccine-escape mutant? By placing the two strains on a phylogeny of other sequenced *H. influezae* genomes, we found that the two strains were unrelated. This surprising result led us to ask a second question:
2. Do these two unrelated strains share particular genes that might have allowed them both to emerge at the same place and time?

## METHODS

### Strain collection and patient characteristics

*H. influenzae* strains were isolated from blood culture of two unrelated individuals (Table 1), both children under five years of age. They presented to Rady Children’s Hospital within one week of each other in January 2016. They were both treated with antibiotics and were eventually cured with no apparent complications. Blood culture was *H. influenzae* positive for both patients and showed that these strains were non-type b, but with an encapsulated appearance. Both strains were sent to the United States Centers for Disease Control and Prevention (CDC) for serotyping, which confirmed them both to be type a. Further patient characteristics are given in Table 1. Strains isolated from patients 1 and 2 were respectively named H1 and H2 in this study.

**Table 1.** Patient characteristics.

|  | Patient #1 | Patient #2 |
| --- | --- | --- |
| Positive culture | Blood culture | Blood and Cerebrospinal fluid (CSF) culture |
| CSF cell profile on first tap | Protein 267<br>Glucose 47<br>Erythrocytes 112,<br>Nucleated cells 619 | Protein 68<br>Glucose 49<br>Erythrocytes 28<br>Nucleated cells 707 |
| CNS complications | Subdural empyema | Seizure |
| Antibiotic Treatment | Vancomycin and meropenem for 18 days | Ceftriaxone for 10 days |
| Time to fever resolution | 14 days | 1 day |

### Ethical approval

This study was approved by the Internal Ethical Review Board and the Privacy Board of Rady Children’s Hospital-San Diego (file #19005C). The study was deemed to be a case report, which does not involve a systematic investigation and therefore does not meet the definition of research as outlined in 45 CFR 46.102(d) and are not subject to the Human Subject Regulations (45 CFR 46). The privacy board concluded that no protected health information (PHI) is disclosed in this study.

### DNA extraction & sequencing

*Haemophilus influenzae* strains were cultured overnight on chocolate agar plates (Thermo Fischer Sceintific) and DNA was extracted using the Bactozol DNA extraction kit (MRC inc.). Extracted DNA was further purified using the PowerClean^®^ Pro DNA Clean-Up Kit (MOBIO Laboratories Inc.). Libraries were prepared using the Nextera XT kit (Illumina Inc.) following the standard Illumina protocol and library size was confirmed at approximately 1000 bp with a Qiaxcel Advanced System (QIAGEN). We performed paired end sequencing (2 x 300 bp) using the MiSeq reagent Kit V3 (Illumina Inc.) on the MiSeq system (Illumina Inc.) yielding a total of 1,128,523 paired-end reads for H1 and 1,708, 296 paired-end reads for H2.

### Genome assembly, annotation, and phylogenetics

Sequencing reads were trimmed with Trimmomatic (Bolger et al., 2014) with default parameters. Trimmed reads were assembled into contigs using IDBA (Peng et al., 2012). We then compared the H1 and H2 genomes with a dataset of 80 non-typable, six b-capsule and one f-capsule genomes (previously described by De Chiara et al. 2014) using gene-by-gene alignments, described below. Consistent with the previous analysis of De Chiara et al., we identified six well-supported clades (named I-VI, following their nomenclature). To ensure that we did not miss close relatives of H1 or H2 not present in the De Chiara dataset, we searched NCBI for closely related genomes using a set of universal single copy genes extracted from H1 and H2 (Creevey et al., 2011). Using this approach, we identified two additional recently sequenced closely related encapsulated genomes: one f-capsule type isolated in Sweden in 2011 (Resman et al., 2011; Su et al., 2013) and one a-capsule type isolated in Canada in 2011 (NCBI accession number: CP017811.1). Ten sequences of *Haemophilus haemolyticus* were used as an outgroup for phylogenetic analyses. Contigs were annotated using the RAST server (Aziz et al., 2008). Translated gene predicitions were assigned to orthologous groups using Orthofinder (Emms and Kelly, 2015). 941 genes assigned to the core genome (present in single copy in each genome) were aligned using MUSCLE (Edgar, 2004). Concatenation of these aligned genes was to infer a core genome phylogeny using FastTreeMP (Price et al., 2009). We also used FastTreeMP to reconstruct each individual gene tree (or gene fragment for genes in the capsule operon). Phylogenies were displayed in FigTree (http://tree.bio.ed.ac.uk/software/figtree/). Gene presence/absence heatmaps were displayed in R with the heatmap.2 function of the ggplots package (http://www.R-project.org/). The same exact protocole of alignment and phylogenetic inference was applied on the smaller regions of the capsule genes.

### Horizontal gene transfer detection

To infer putative recent recombination due to horizontal gene transfer (HGT) between two strains, we used an explicit phylogenetic method where individual gene trees (including both core and flexible genes) were screened for phylogenetic incongruence with the reference tree, containing six major lineages, based on 941 core genes. A phylogenetic incongruence was considered as an HGT if a gene sequence from a lineage [i] was grouped in a well-supported clade (bootstrap value of 100) with a gene from a different lineage [j]. We also required the gene sequences from lineages [i] and [j] to share >97% nucleotide identity. As some flexible genes are shared uniquely between two isolates from different clades, we also considered these as putative HGTs if they shared >97% nucleotide identity.

## RESULTS

### H1 and H2 genomes are distantly related

The core genome phylogeny based on 941 aligned genes shows that H1 and H2 belong to two distinct lineages: H1 in clade VI and H2 in clade I (**Figure 1**). These two clades are distantly related, clade I being the first *H. influenzae* branch after its divergence from the *H. haemolyticus* outgroup. Within clade VI, H1 is the only type-a isolate in a sub-clade carrying b-type capsule genes, with the exception of one genome, Hi1008, which likely lost most of its capsule genes (Meyler et al., 2019). We searched the NCBI genome database to identify close relatives of H1 and H2 not present in our phylogeny. The closest sequenced relative of H1, NML_Hia1, was isolated in Canada in 2011, and also carries an a-type capsule locus. The closest sequenced relatives of H2 are two f-type genomes: Hi794 (Finland, date unknown) and KR494 (Sweden, 2011). As H1 and H2 are distantly related in the tree, they are clearly not epidemiologically linked and likely represent independent infection events.

**Figure 1.**
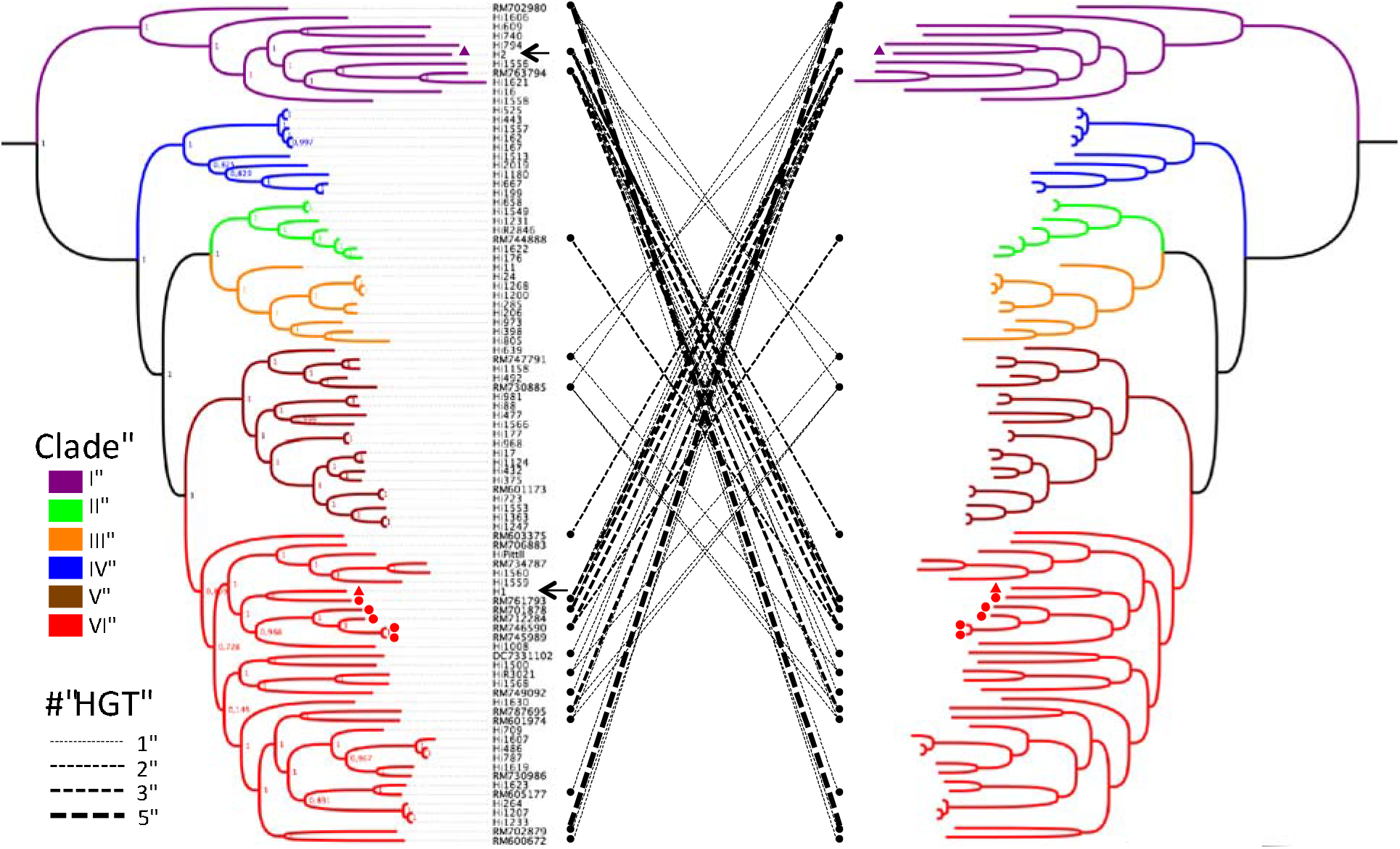
Core genome phylogeny of *H. influenzae* and putative HGT events between clades. *H. haemolitycus* was used as an outgroup to root the tree (not shown). Encapsulated strains are indicated with a circle (b-type) or a triangle (a-type); the remaining strains are unencapsulated. H1 and H2 are indicated with arrows. Clades I-VI (following the nomenclature of De Chiara et al. 2014) are indicated in different colours. Putative HGT events between clades are indicated with dashed lines. Line thickness indicates the number of HGTs (ranging from 1 to 5 genes).

### H1 and H2 belong to clades engaging in pervasive HGT

Despite being phylogenetically unrelated, H1 and H2 were both isolated during the same week from the same hospital. We thus hypothesized that they may have recently exchanged genes via HGT. Using phylogenetic criteria to detect HGT (Methods) we did not identify any recent gene transfers between H1 and H2 (**Figure 1**). Rather, H1 and H2 had flexible gene profiles similar to other genomes from their respective clades (**Figure S1**). Despite the lack of recent HGT between H1 and H2, we did observe that HGTs were unevenly distributed between clades (chi-square = 163.46, df = 14, *P* < 2.2e-16) and that the two distantly related clades containing H1 and H2 (clades I and VI) engaged in particularly pervasive HGT (**Figure 1**). These HGTs encode a mix of virus- and plasmid-related functions, antibiotic resistance genes, and metabolic genes (**Table S1**). Clades I and VI have similar profiles of selected virulence factors (De Chiara et al. 2014), and members of both clades tend to encode *hap*, *hia*, and *hif* virulence genes (**Figure S2**). Neither clade has a strong preference for a particular geographic region (**Figure S3**), time period (**Figure S4**), or infection site (**Figure S5**). It is thus unclear why clades I and VI apparently engage in more frequent HGT than other clades.

### Detailed phylogeny and HGT of the capsule locus

The presence of two a-type strains (H1 and H2) in two distantly-related clades lead us to investigate in greater details the evolution of capsule locus genes. By manually inspecting individual gene alignments at this locus, we found that H1 and H2 had identical or very similar sequences spanning a ~5kb region encompassing most of the serotype-specific genes (**Figure 2**). These genetic similarities between H1 and H2 were not detected in the gene-by-gene analysis (**Figure 1**) because of conflicting phylogenetic signals within genes (**Figure 2**). NML_Hia1, another a-type genome isolated in Canada in 2011, also shared a similar or identical sequence in the serotype-specific region, suggesting that the putative recombination event in this region occurred in 2011 or earlier, and that the sequence has subsequently accumulated relatively few mutations. Upstream and downstream of the serotype-specific region, H2 was most similar to f-type strains (**Figure 2**). This suggests that an f-type ancestor of H2 acquired ~5kb of serotype-specific DNA from an a-type donor strain, resulting in a serotype switch. Thus, recent ancestors of H1 and H2 engaged in HGT at the capsule locus. However, the H1 and H2 are non-identical, notably in the *acsC* gene (6 substitutions over 2 400 bp) (**Figure 2**), making it unlikely that the exchange occured in Rady Children’s Hospital.

**Figure 2:**
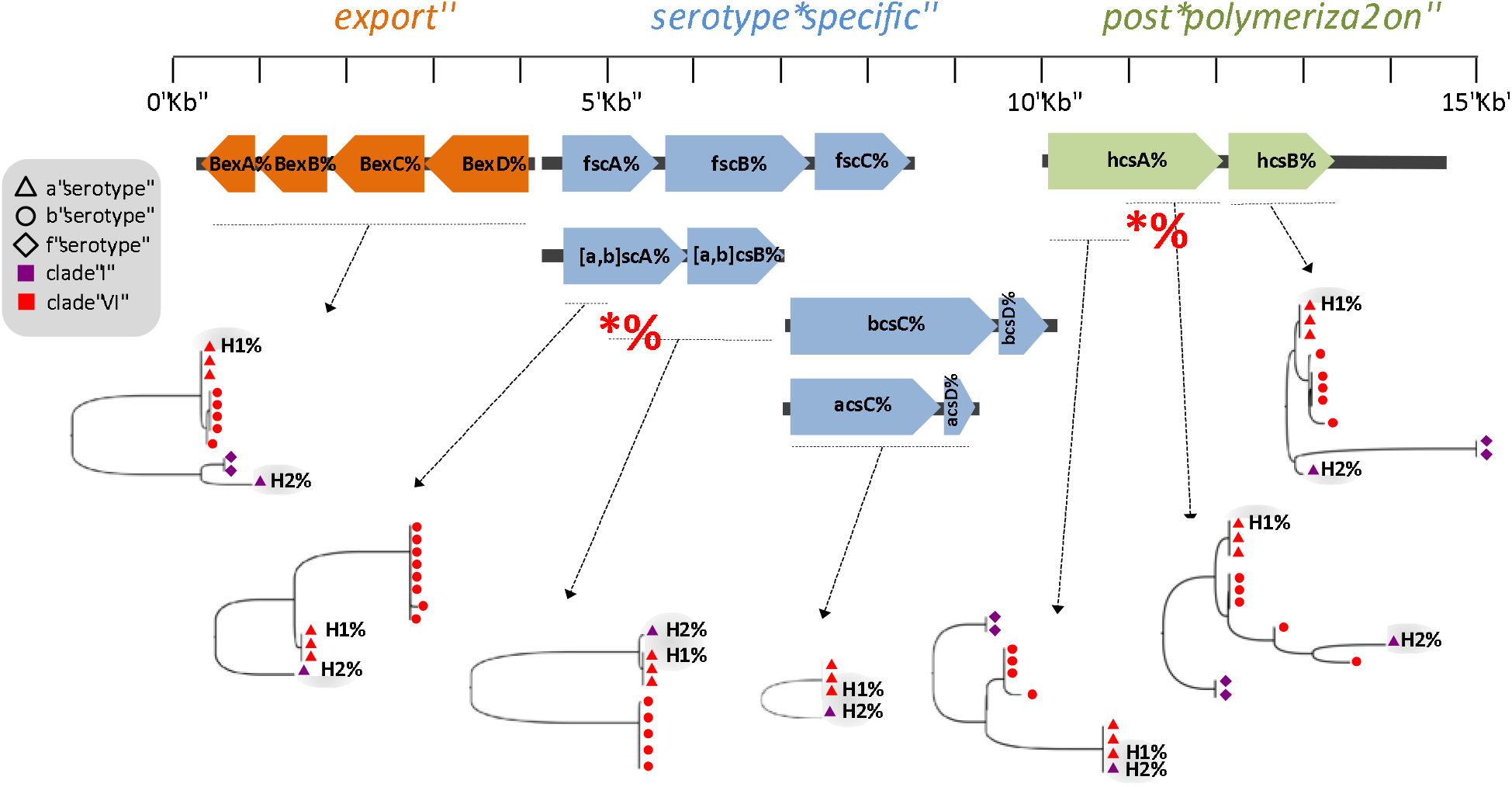
Phylogenies of genes and regions within the capsule locus. Dashed horizontal lines indicate the region used for phylogenetic analysis. Red stars indicate putative recombination breakpoints.

## DISCUSSION

The appearance of two *H. influenzae* infections in the same hospital in the same week was highly unexpected because such infections are exceedingly rare in areas of high vaccine coverage. This raised concerns that the two cases were part of a local outbreak, involving *H. influenzae* transmission in the San Diego area. By sequencing the two isolate genomes (H1 and H2) and placing them on a phylogenetic tree encompassing the known diversity of *H. influenzae*, we found that they belonged to distantly related clades, indicating that each infection was aquired independently and the two were not linked in a recent transmission chain. The entire analysis, from strain isolate to sequencing and phylogenetic analysis, could be performed in about a week, allowing us to rapidly rule out local transmission. Rather, the observation that H1 and H2 are distantly related highlights the potential for rapid, potentially global spread of different *H. influenzae* lineages.

To our knowledge H2 is the first North American clade I genome to be sequenced, suggesting a recent transmission from Europe or Asia (**Figure S3**). H1 is part of clade VI and shares recent ancestry with another type-a Canadian isolate from 2011, suggesting that this type-a lineage has been circulating in North America for several years. Despite being distantly related, H1 and H2 share a very similar capsule locus, particularly in the serotype-specific region. Flanking the serotype-specific region and elsewhere in the genome, H2 is more similar to f-type strains. Together, these observations point to a scenario in which an f-type ancestor of H2 arrived in North America around 2011 and acquired an a-type capsule by recombination with a recent ancestor of H1. We also note that H1 and H2 belong to two clades (VI and I, respectively) that appear to engage in preferential HGT, possibly to cryptic niche overlap. Future work will be needed to confirm and understand the reasons for this preferential genetic exchange.

That both H1 and H2 were isolated at the same place and time appears to be a coincidence, but does suggest that these a-type strains are filling a vacant niche left by b-type strains targeted by current vaccines. Our results also indicate that vaccines need not select for lineage replacement, but could allow multiple different lineages to adapt via acquisition of new capsule loci by HGT. We show that such a scenario is plausible, but further analysis of larger population genomic samples will be needed to assess the relative importance of lineage replacement vs. capsule HGT in the evolutionary response of *H. influenzae* to vaccine pressure.

## Supplementary Table

**Table S1.**
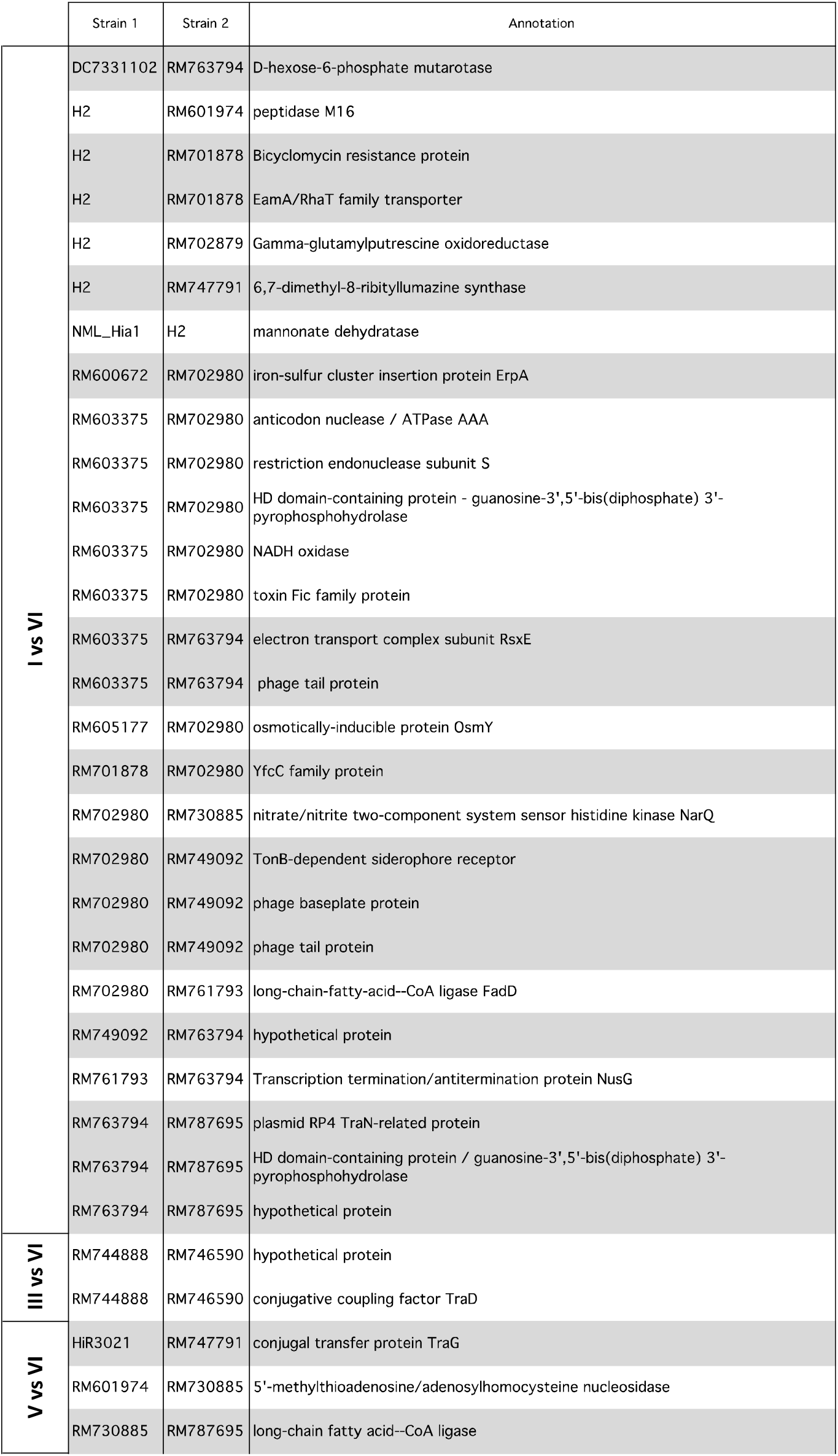
Annotated functions of putative HGTs between clades. Strain 1 and 2 indicated the two strains involved in the putative HGT. Clades involved (I-VI) are shown in the far left column.

