## Supplementary Figures for "Two cases of type-a *Haemophilus influenzae* meningitis within the same week in the same hospital are phylogenetically unrelated but recently exchanged capsule genes"

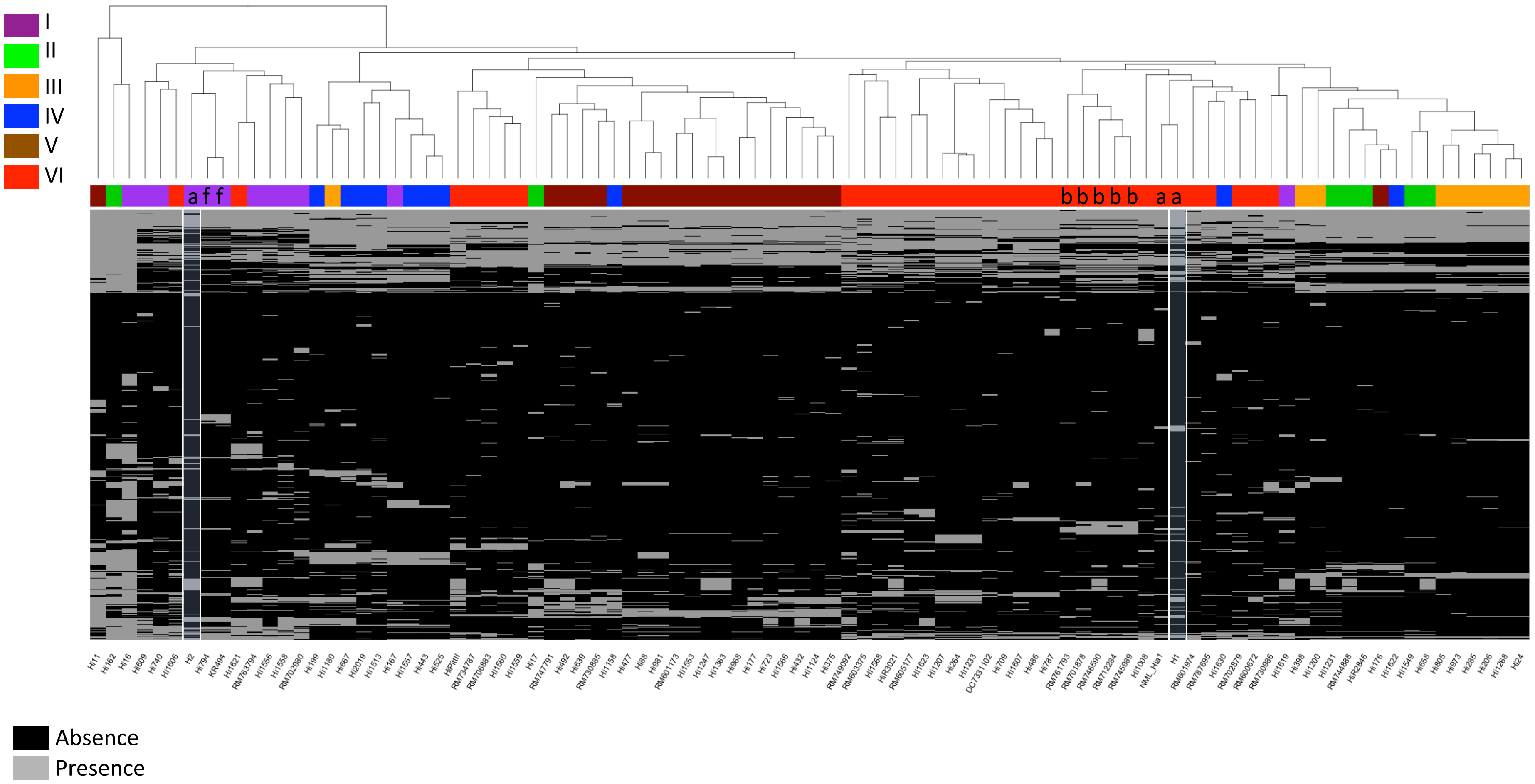


**Figure S1** : Clustering of genomes according to their flexible gene content.

Encapsulated strains are indicated with letters (a,b, or f) at the top of the heatmap. Clades I-VI are indicated in different colours.

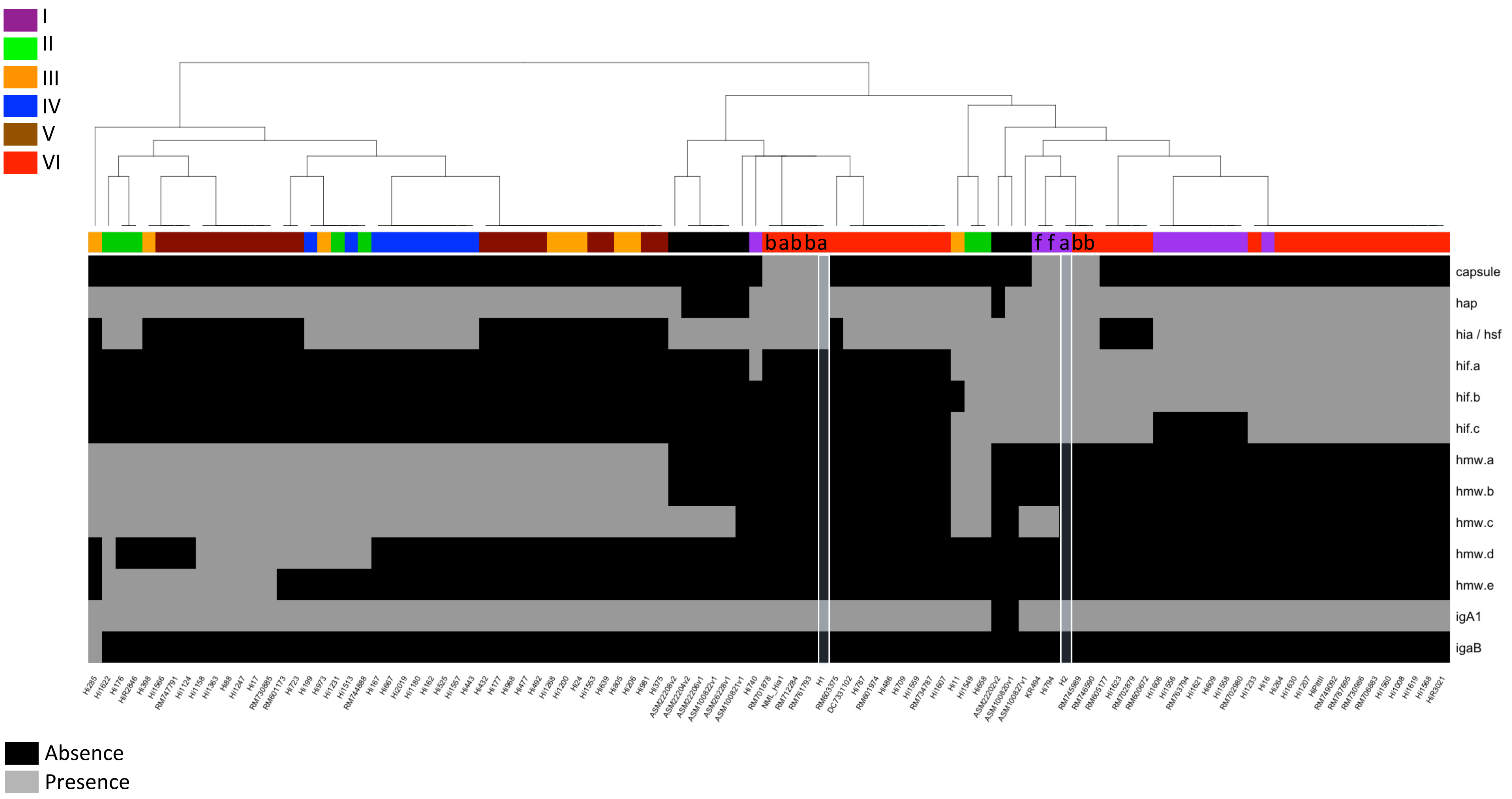


**Figure S2 :** Profile of selected virulence gene content across genomes.

Strains are clustered according to their virulence gene content. H1 and H2 are indicated with white boxes. Encapsulated strains are indicated with letters (a,b, or f) at the top of the heatmap. Clades I-VI are indicated in different colours.

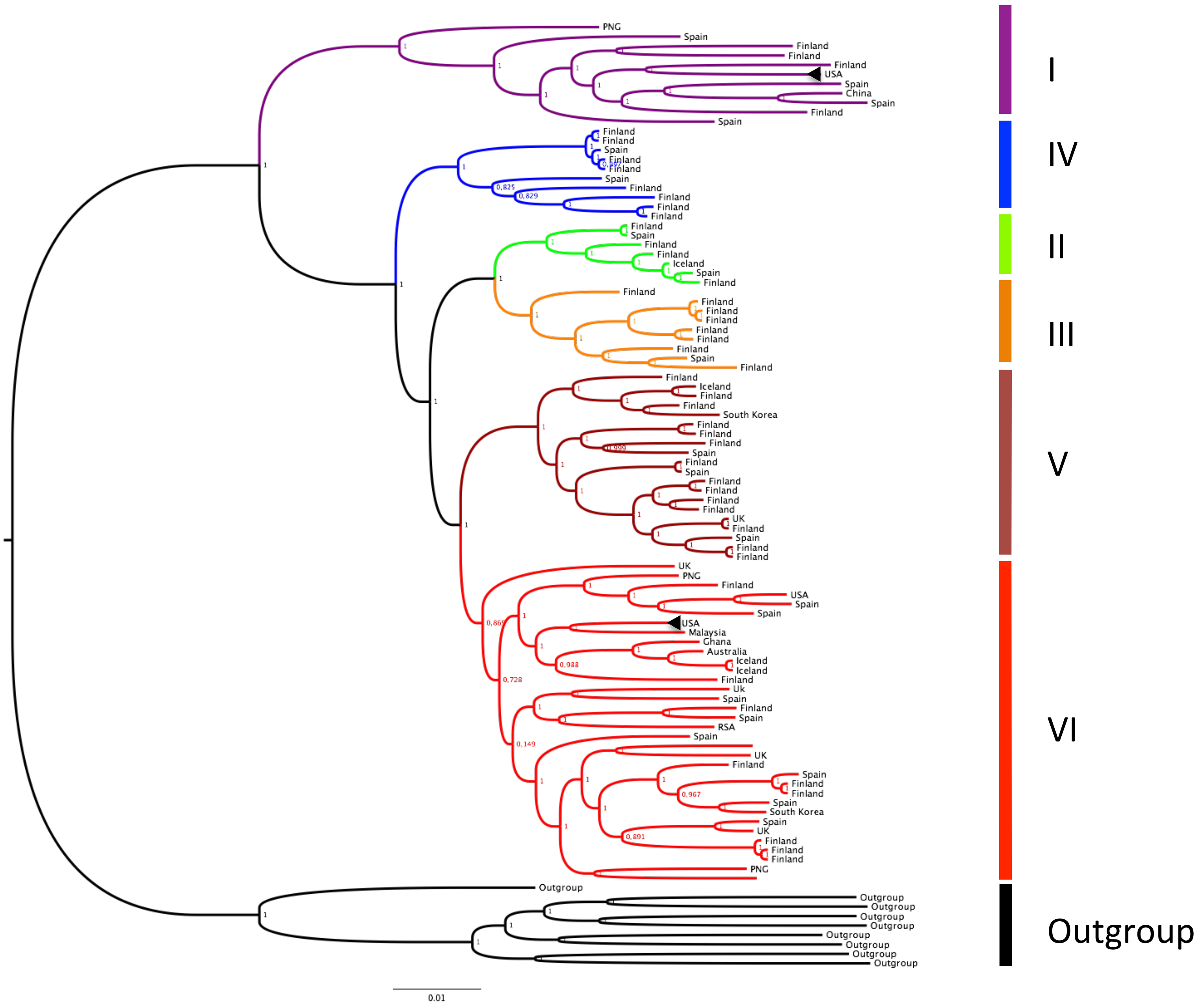


**Figure S3**: Core genome phylogeny of *H.influenzae* annotated by country of origin.

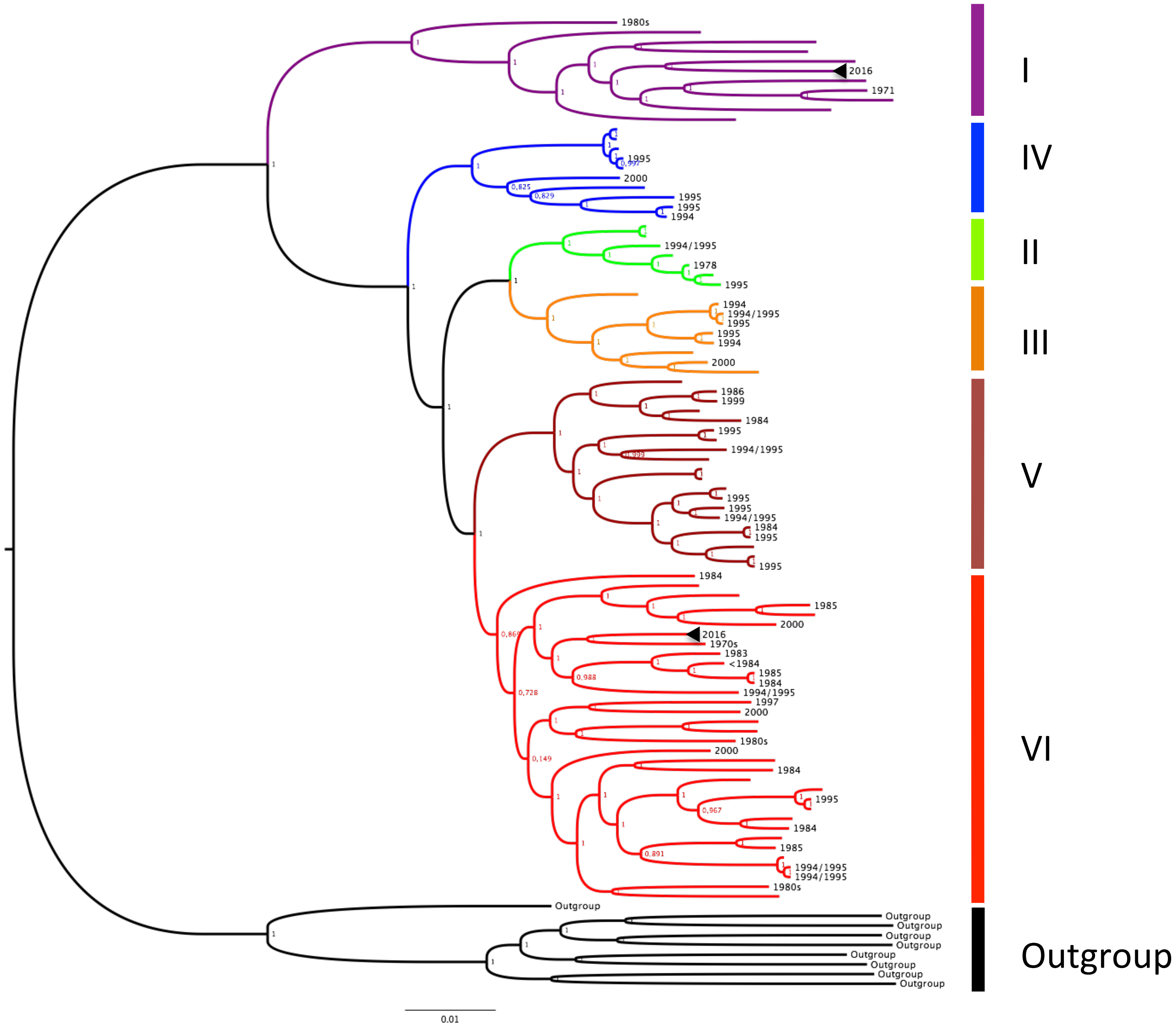


**Figure S4**: Core genome phylogeny of *H.influenzae* annotated by year of sampling.

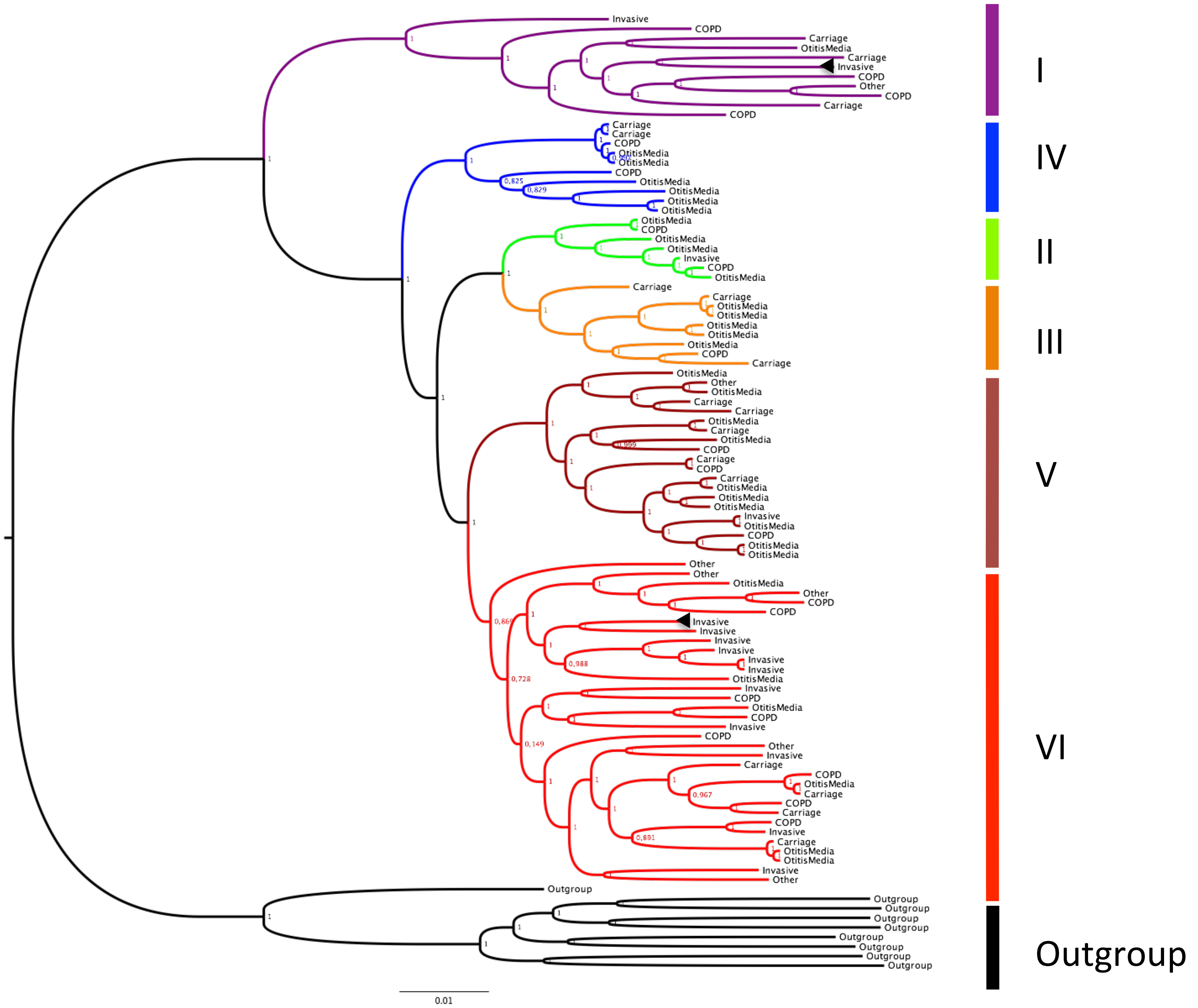


**Figure S5**: Core genome phylogeny of *H.influenzae* strains annotated by disease type
